# Pan-cancer Proteomics Analysis to Identify Tumor-Enriched and Highly Expressed Cell Surface Antigens as Potential Targets for Cancer Therapeutics

**DOI:** 10.1101/2023.01.23.525265

**Authors:** Jixin Wang, Wen Yu, Rachel D’Anna, Anna Przybyla, Matt Wilson, Matthew Sung, John Bullen, Elaine Hurt, Gina DAngelo, Ben Sidders, Zhongwu Lai, Wenyan Zhong

**Author notes:** **Corresponding author:** Wenyan Zhong, AstraZeneca, One MedImmune Way, Gaithersburg, MD 20878, USA.

## Abstract

The National Cancer Institute’s Clinical Proteomic Tumor Analysis Consortium (CPTAC) provides unique opportunities for cancer target discovery using protein expression. Proteomics data from CPTAC tumor types have been primarily generated using a multiplex tandem mass tag (TMT) approach, which is designed to provide protein quantification relative to reference samples. However, relative protein expression data is suboptimal for prioritization of targets within a tissue type, which requires additional reprocessing of the original proteomics data to derive absolute quantitation estimation. We evaluated the feasibility of using differential protein analysis coupled with intensity-based absolute quantification (iBAQ) to identify tumor-enriched and highly expressed cell surface antigens, employing tandem mass tag (TMT) proteomics data from CPTAC. Absolute quantification derived from TMT proteomics data was highly correlated with that of label-free proteomics data from the CPTAC colon adenocarcinoma cohort, which contains proteomics data measured by both approaches. We validated the TMT-iBAQ approach by comparing the iBAQ value to the receptor density value of HER2 and TROP2 measured by flow cytometry in about 30 selected breast and lung cancer cell lines from the Cancer Cell Line Encyclopedia. Collections of these tumor-enriched and highly expressed cell surface antigens could serve as a valuable resource for the development of cancer therapeutics, including antibody-drug conjugates and immunotherapeutic agents.

## INTRODUCTION

Biologics-based cancer therapeutics including antibody drug conjugates and immunotherapies are proven modalities for the treatment of cancer. Most of these therapeutics often target proteins preferentially expressed on the surface of cancer cells (1). For many years, drug candidate targets for these modalities have been discovered through mining of large RNA expression databases such as The Cancer Genome Atlas (TCGA) as a surrogate approach in the absence of a large protein expression database (2). Recent advancements in proteogenomic characterization of large collections of cancer patient samples represented by The National Cancer Institute’s Clinical Proteomic Tumor Analysis Consortium (CPTAC) (3) has provided unique opportunities for cancer target discovery using protein expression.

The CPTAC serves as a repository of global proteomics data of tumor and normal adjacent tissue (NAT) samples from more than 10 tumor types, including colon adenocarcinoma (COAD) (4), breast cancer (BRCA) (5), lung adenocarcinoma (LUAD) (6), ovarian cancer (OV) (7), glioblastoma multiforme (GBM) (8), clear-cell renal cell carcinoma (ccRCC) (9), pancreatic ductal adenocarcinoma (PDA) (10), uterine corpus endometrial carcinoma (UCEC) (11), head and neck squamous-cell carcinoma (HNSCC) (12), and lung squamous-cell carcinoma (LSCC) (13). However, proteomics data from these tumor types have been primarily generated by the multiplex tandem mass tag (TMT) approach, which was designed to provide relative protein quantification to reference samples. In addition to preferential expression in tumor compared to normal tissues, ideal targets for biological therapeutics often requires high level expression of these targets on the tumor cell surface. The relative protein expression data from CPTAC hinders the direct utilization of the proteomics data for target prioritization and additional reprocessing of the original proteomics data to derive absolute quantitation estimation is required.

Mass spectrometry–based proteomics methods have been widely used for relative and absolute quantification of proteomes and posttranslational modifications (14). Both label-free and label-based quantification methods have been applied in proteomics studies. Several algorithms for label-free absolute protein quantification have been developed, including TOP3 (15), label-free absolute quantification (LFAQ) (16), absolute protein expression (APEX) (17), total protein approach (TPA) (18–21), and intensity-based absolute quantification (iBAQ) (22). The TPA approach has been applied to the data-independent acquisition (DIA)–based label-free quantification (LFQ) method, and the tandem mass spectrometry–based DIA-TPA approach is comparable to the mass spectrometry–based data-dependent acquisition (DDA)–TPA method (23).

In addition to these methods, FragPipe (24) with the MSFragger search engine can provide equivalent protein intensity data based on the intensities of the precursors and TMT reporters, which can be used as input to derive estimated absolute quantitation by using previously published absolute quantification algorithms. This approach has previously been applied to CPTAC ccRCC data processing (9). Although the intensity calculated by FragPipe reflects the absolute intensity of the protein summarized from intensities from all ionized peptides. However, this value cannot be used to directly compare the protein abundance across different proteins, since different proteins have different number of ionizable peptides. In this study, we developed a computational approach to estimate absolute protein abundance from TMT proteomics data. We reprocessed TMT proteomics data with FragPipe (25) and calculated the TMT-TPA and TMT-iBAQ values for 10 tumor types. Two methods for calculating estimated absolute protein abundance were compared using label-free global proteomics data from the CPTAC COAD data set, which contains both label-free and TMT proteomics data. We also compared the LFQ-TPA (label-free) and the TMT-TPA (TMT) value of the CPTAC COAD data set. The iBAQ approach was evaluated by using receptor density values (26) measured by flow cytometry and assessing the potential application of the iBAQ data for comparing protein expression levels by indication to identify tumor-enriched and highly expressed cell surface antigens. Finally, by combining differential analysis of tumor and NAT samples with surface protein prediction, we were able to identify pan-cancer tumor-enriched and highly expressed proteins as potential cancer antigens for target discovery (27, 28), cancer vaccine development (29), and cancer biology studies (30).

## EXPERIMENTAL SECTION

### Re-analysis of CPTAC and CCLE data sets

Raw data files from the CPTAC were downloaded from https://proteomic.datacommons.cancer.gov/pdc/ with PDC study identifier COAD (PDC000116), BRCA (PDC000120), ccRCC (PDC000127), PDA (PDC000393), OV (PDC000118), LSCC (PDC000234), GBM, LUAD (PDC000153), HNSCC (PDC000221), and UCEC (PDC000125), and data files from the Cancer Cell Line Encyclopedia (CCLE) were downloaded from https://massive.ucsd.edu/ProteoSAFe/dataset.jsp?task=02cd1b6a7c674f3ebdbed300b5d9aa57 (MassIVE repository accession MSV000085836). The CPTAC COAD data set contains both label-free and TMT proteomics data. LFQ was carried out on the unlabeled data (100 tumor samples) of COAD from CPTAC, using MaxQuant (31). TMT 10-plex mzML files from CPTAC COAD, BRCA, ccRCC, PDA, OV, LSCC, GBM, LUAD, HNSCC, and UCEC and CCLE were analyzed via the FragPipe computational platform (version 15) with MSFragger (version 3.2) and Philosopher (version 3.4.13) (25) for protein identification and quantitation, using the recommended search parameters with global normalization. Protein grouping was performed.

The label-free data were searched in MaxQuant against a collection of 73045 protein sequences downloaded from Uniprot on June 22, 2018, with full tryptic specificity and up to two miss-cleavage sites, fixed modification of “Carbamidomethyl (C)”, and variable modifications of “Oxidation (M), Acetyl (Protein N-term) and Deamidation (NQ)”. The mass tolerance was set to 20 ppm for both the precursors and the fragments. The protein identifications were filtered at the 1% false discovery rate (FDR) level as determined by MaxQuant.

The TMT-labelled data, they were searched in FragPipe (version 15.0 with MsFragger version 3.2 and Philosopher version 3.4.13) against protein sequences (reviewed, 20420 entries) downloaded from Uniprot by FragPipe on July 15, 2021, with full tryptic specificity and up to two miss cleavages, fixed modification of Cys residue by 57.02146 daltons (Da), and Lys residue by 229.16293 Da, variable modifications of Met residue by 15.9949 Da, protein N-terminus by 42.0106 Da, and peptide N-terminus or Ser residue by 229.16293 Da. The mass tolerance was set to 20 ppm for both the precursors and the fragments. The protein identifications were filtered to 1% FDR as determined by FragPipe from the embedded decoy sequences.

LFQ-TPA and LFQ-iBAQ values of the CPTAC COAD cohort were calculated from label-free raw intensity using equations 1 and 2 (see below) respectively. TMT-TPA, TMT-iBAQ, and TMT-iBAQ-derived copy number values of all 10 CPTAC indications and CCLE were calculated from protein abundance output of FragPipe using equations 1, 2, and 3 (see below), respectively.

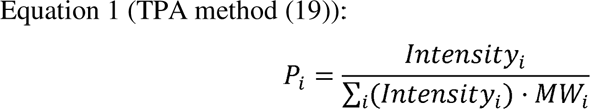

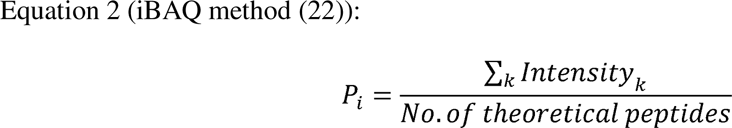

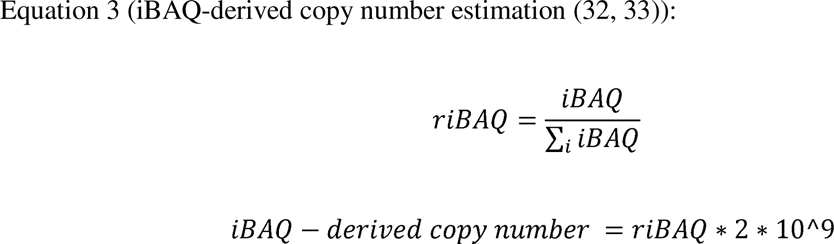

where *Pi* = predicted absolute abundance of protein *i,* and *k* = peptide and MW = molecular weight. TPA units are expressed as femtomoles per gram. “No. of theoretical peptides” refers to the number of theoretical peptides when that protein is typically digested by trypsin. iBAQ values were calculated based on UniProt protein accessions and reported using associated gene symbols. For iBAQ-derived copy number estimation, 2 × 10^9^ as the total number of proteins in a cell was used based on published studies (33).)

### Differential protein analysis

FragPipe protein abundance data from CPTAC tumor and NAT samples were used for differential protein analysis. Principal component analysis and unsupervised hierarchical clustering were used to evaluate batch effect and sample quality control. An initial filtering step was applied to the protein abundance data, and proteins that were quantified in at least 50% of the samples were used for further analysis. Missing value visualization was performed, and k–nearest neighbor (KNN) imputation was applied to the data with the R package MSnbase (version 2.20.4) (34). Previous studies have shown that KNN imputation performs better in TMT proteomics data (35, 36). Differential protein expression analysis was performed with R (version 4.1.1), using Empirical Bayes statistics on protein-wise linear models with limma (37) embedded in the differential protein analysis (DEP) package (version 1.16.0) (38). Proteins that were significantly upregulated in tumor compared with NAT were identified with a threshold of adjusted *P* = 0.01 and fold change = 1.5. Cell surface proteins were predicted using surface prediction consensus (SPC) score downloaded from SurfaceGenie (39–41), and proteins with SPC ≥ 1 were annotated as potential surface proteins. Differentially upregulated cell surface proteins were ranked by their iBAQ values in tumor tissues to obtain highly expressed, tumor-enriched cell surface antigens.

### Flow experiment and analysis methodology

Datopotamab and trastuzumab deruxtecan (T-DXd) (human immunoglobulin G1 [IgG1]) were labeled with Alexa Fluor 647, using the Alexa Fluor protein labeling kit (A20173; Invitrogen). Cells from CCLE cell lines were harvested with Accutase (A1110501; Gibco) and stained with Live/Dead Fixable Violet Dead Cell Stain (L34955; Invitrogen) for 20 minutes on ice and protected from light. Cells were then washed and stained with 20 μg of datopotamab–Alexa Fluor 647 or T-DXd–Alexa Fluor 647 per mL for 1 hour on ice and protected from light. Cells were fixed with 2% paraformaldehyde (AAJ19943K2; Thermo Fisher Scientific) for 20 minutes at room temperature and stored in phosphate-buffered saline (PBS) at 4°C overnight.

Quantum Simply Cellular anti-human IgG beads (816; Bangs Laboratories) were stained with 20 μg of datopotamab–Alexa Fluor 647 or T-DXd–Alexa Fluor 647 in PBS for 30 minutes at room temperature. Beads were then washed three times in PBS and transferred to flow tubes.

Beads and cells were run on an LSRFortessa flow cytometer (BD Biosciences), using the same voltage settings. Flow data analysis was performed on an FCS Express flow cytometer (De Novo Software) and bead median fluorescence intensity values were used to plot the ABC standard curves for datopotamab and T-DXd. The lot-specific template provided with the bead kit was used to obtain cell ABC values by extrapolating from the standard curves.

### Correlation analysis

Correlations of FragPipe protein abundance with global normalization and LFQ intensity, iBAQ vs. TPA, LFQ-TPA vs. TMT-TPA, and LFQ-iBAQ vs. TMT-iBAQ, as well as receptor density vs. iBAQ, and protein abundance by FragPipe vs derived iBAQ data were determined with Pearson correlation.

## RESULTS

### Differentially up-regulated proteins in tumor compared to NAT

We first identified proteins that were upregulated in tumor tissues compared with NAT. The workflow used in this study is outlined in Figure 1. We re-analyzed the TMT 10-plex mzML files of 10 indications (COAD, BRCA, LUAD, OV, GBM, ccRCC, PDA, UCEC, HNSCC, and LSCC) from CPTAC through FragPipe and MSFragger to obtain protein abundance values for identified proteins expressed in tumor and NAT. The protein summary and quantification from FragPipe and MSFragger output for CPTAC 10 indications are shown in Supplemental Table S1. Missing values are commonly observed in proteomics data due to a variety of reasons, including protein digestion loss, measurement failure, protein abundances below the instrument limit of detection, poor ionization efficiency, and signal-to-noise ratio (42–44). It is critical to apply the correct missing value imputation method before performing differential abundance analysis. Our evaluation of the missing values showed a missing-at-random pattern across tumor and normal conditions but a missing-not-at-random pattern at the multiplex level. Therefore, we chose a KNN algorithm to impute missing values (45) and identified a range of differentially upregulated proteins across various tumor types (Table 1). As an example, Supplemental Figure S1 shows a missing data pattern for the CPTAC ccRCC indication.

**Figure 1.**
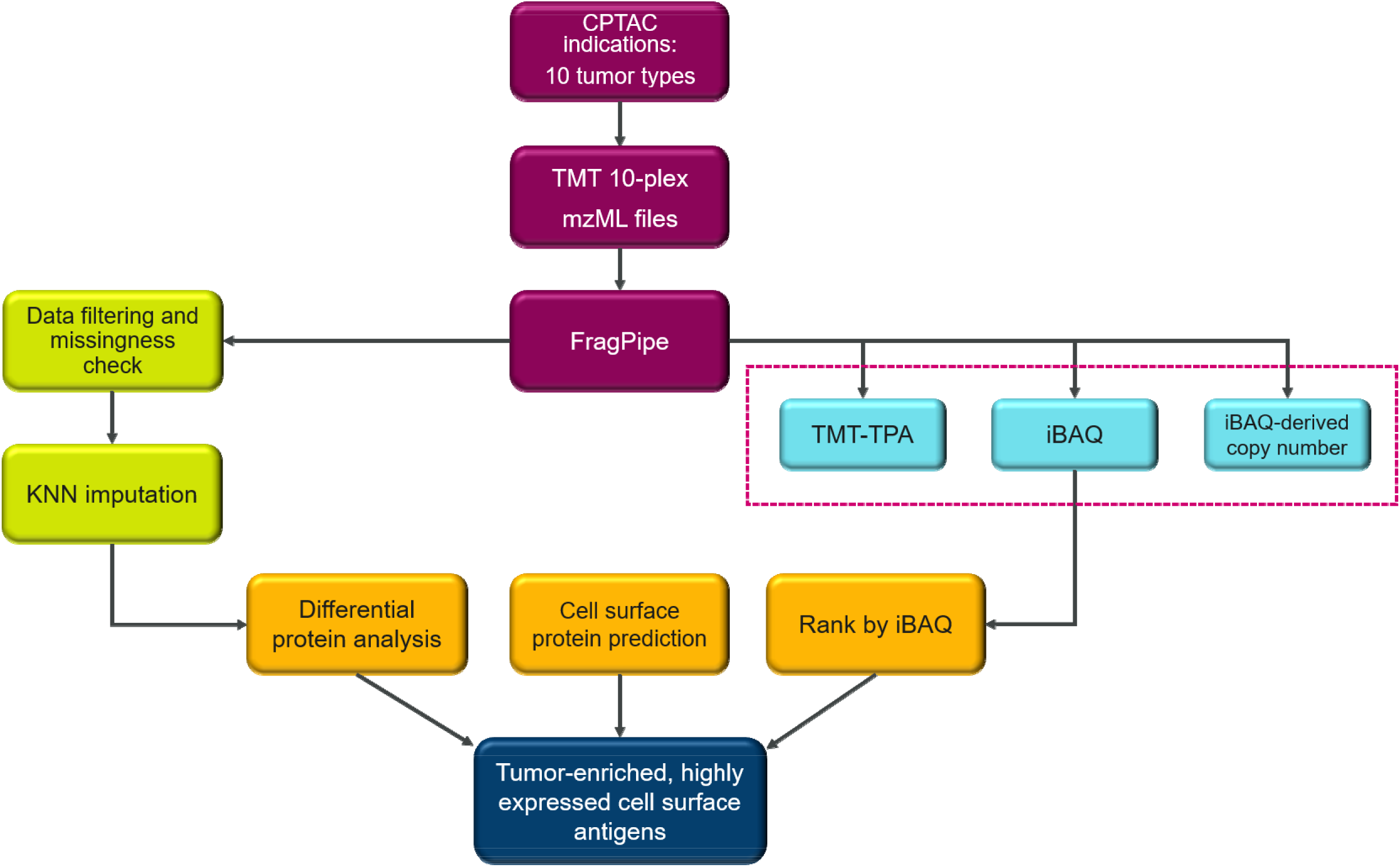
Workflow for identification of tumor-enriched and highly expressed cell surface antigens.

**Table 1.** Summary of differential expression analysis and top-ranked tumor enriched and highly expressed tumor antigens.

| **CPTAC indication** | **No. of upregulated proteins** | **No. of predicted cell surface proteins** | **Top 5 tumor-enriched antigens** |
| --- | --- | --- | --- |
| BRCA | 210 | 64 | VAMP8, PYCARD, TMED2, IER3IP1, DPM3 |
| COAD | 236 | 46 | ELANE, PRTN3, CEACAM5, FCER1G, GPRC5A |
| ccRCC | 968 | 249 | MIF, CRYAB, LGALS1, ANXA4, ANXA2 |
| PDA | 242 | 57 | BGN, ANXA1, CD59, CTSE, THBS1 |
| LUAD | 463 | 115 | HSPA5, HSPD1, SEC61B, PDIA4, SPCS3 |
| UCEC | 998 | 417 | CALR, MT-CO2, CD9, ATP5IF1, COX6C |
| OV | 159 | 28 | CALR, HSPD1, HSP90AB1, C1QBP, PDIA4 |
| HNSCC | 477 | 185 | ELANE, PRTN3, HLA-B, SLC2A1, TAPBP |
| GBM | 207 | 52 | APOA1, ANXA5, ANXA1, ANXA2, TF |
| LSCC | 1065 | 209 | HSP90AB1, HSPA5, SLC25A5, HSPD1, VDAC2 |

### Estimation of absolute protein abundance

In addition to differential protein analysis, we estimated absolute protein abundance using TPA and iBAQ approaches. Both methods have been previously applied and validated in estimating absolute protein abundance with label-free proteomics data (23, 46). The CPTAC COAD data set contains both label-free and TMT global proteomics data from the same tumor samples and thus is an ideal data set with which to compare absolute protein abundance calculated from both platforms. The data source and analysis workflow are shown in Supplemental Figure S2. The correlation between FragPipe protein abundance and label-free quantification intensity is 0.67 as shown in Supplemental Figure S3.

We then compared iBAQ and TPA methods for calculating estimated absolute protein abundance, using label-free and TMT global proteomics data from the COAD data set. For label-free proteomics data, LFQ-TPA was highly correlated with LFQ-iBAQ (*r* = 0.95), as shown in Figure 2A. A similar trend was observed for TMT-TPA and TMT-iBAQ correlation (*r* = 0.98) in the TMT proteomics data, suggesting that the TPA and iBAQ methods are highly comparable. Next, we compared LFQ-TPA (label-free) with TMT-TPA (TMT), and LFQ-iBAQ with TMT-iBAQ, using overlap samples and proteins from the COAD data set. The LFQ-iBAQ and TMT-iBAQ methods of the overlap samples and proteins for the COAD data set were well correlated (*r* = 0.71). TMT-iBAQ greatly underestimated protein abundance compared with LFQ-iBAQ (Figure 2B), although a higher correlation was observed between LFQ-TPA and TMT-TPA (*r* = 0.76). Further, we investigated the correlation between protein abundance reported by FragPipe and derived TMT-iBAQ data and the correlation ranges from 0.85 to 0.88 (R > 0.5) across 10 indications (Supplemental Figure S4). These results suggest that both TPA and iBAQ can be applied to TMT proteomics data to estimate absolute protein abundance. We chose to use iBAQ to rank protein expression for identifying highly expressed cell surface antigens because this method is used in most of the protein copy number estimation methods reported in the literature (47–49).

**Figure 2.**
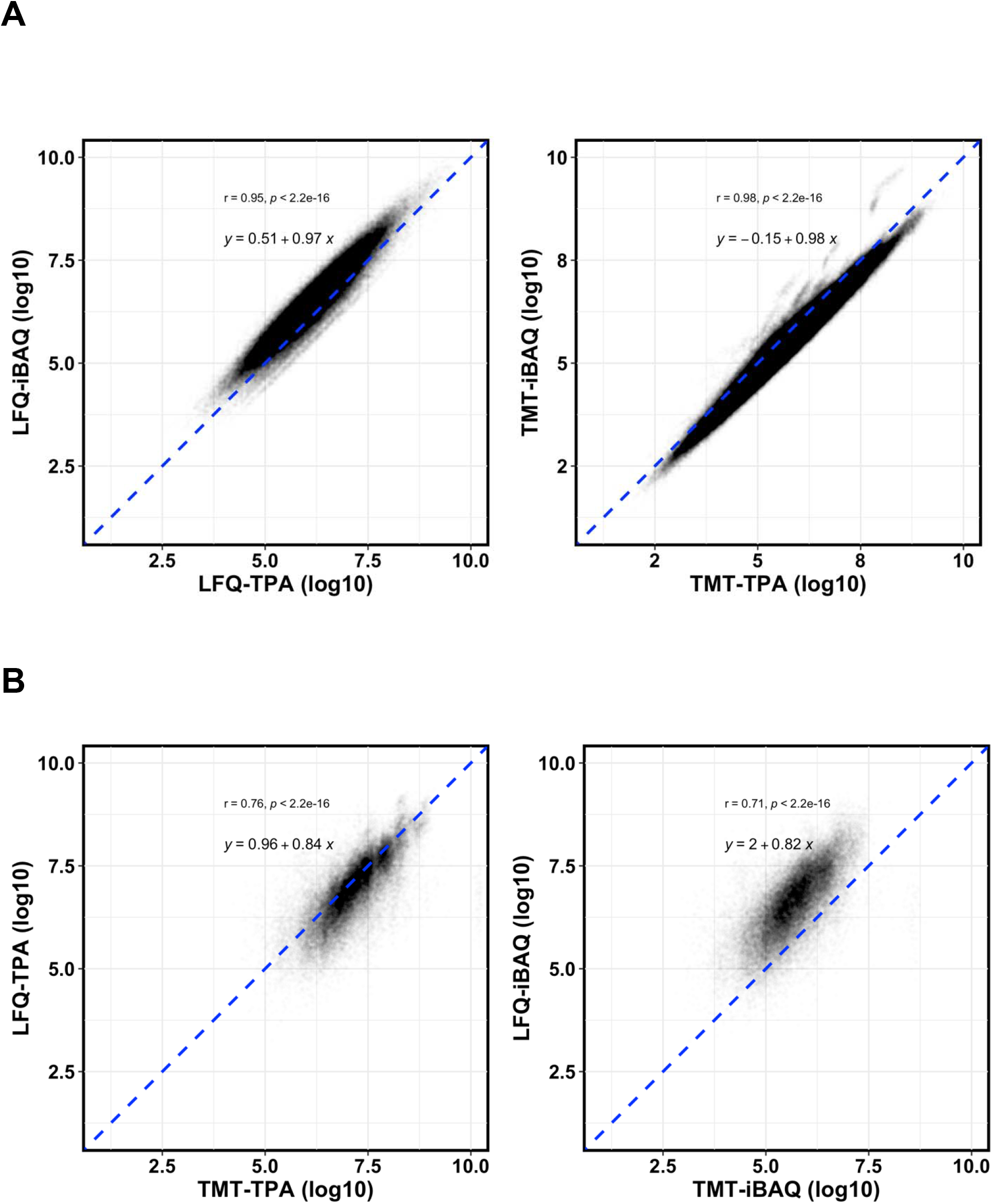
Estimation of absolute protein abundance of iBAQ derived from TMT and label-free proteomics data, respectively, in the CPTAC COAD data set. (A) LFQ-TPA and LFQ-iBAQ, as well as TMT-TPA and TMT-iBAQ, were highly correlated by Pearson correlation. (B). For overlap sample and proteins, the TPA and iBAQ methods showed relatively good correlation between LFQ and TMT data.

### Comparison of TMT-iBAQ absolute protein abundance estimation to receptor density values measured by flow cytometry

To evaluate iBAQ-based protein abundance estimation for TMT proteomics data, we compared experimentally generated receptor density values of HER2 and TROP2 of CCLE cell lines with iBAQ values derived from CCLE TMT proteomics data processed through FragPipe. iBAQ quantity and receptor density for HER2 and TROP2 were highly correlated, with Pearson *r* = 0.82 and *r* = 0.83, respectively (Figure 3).

**Figure 3.**
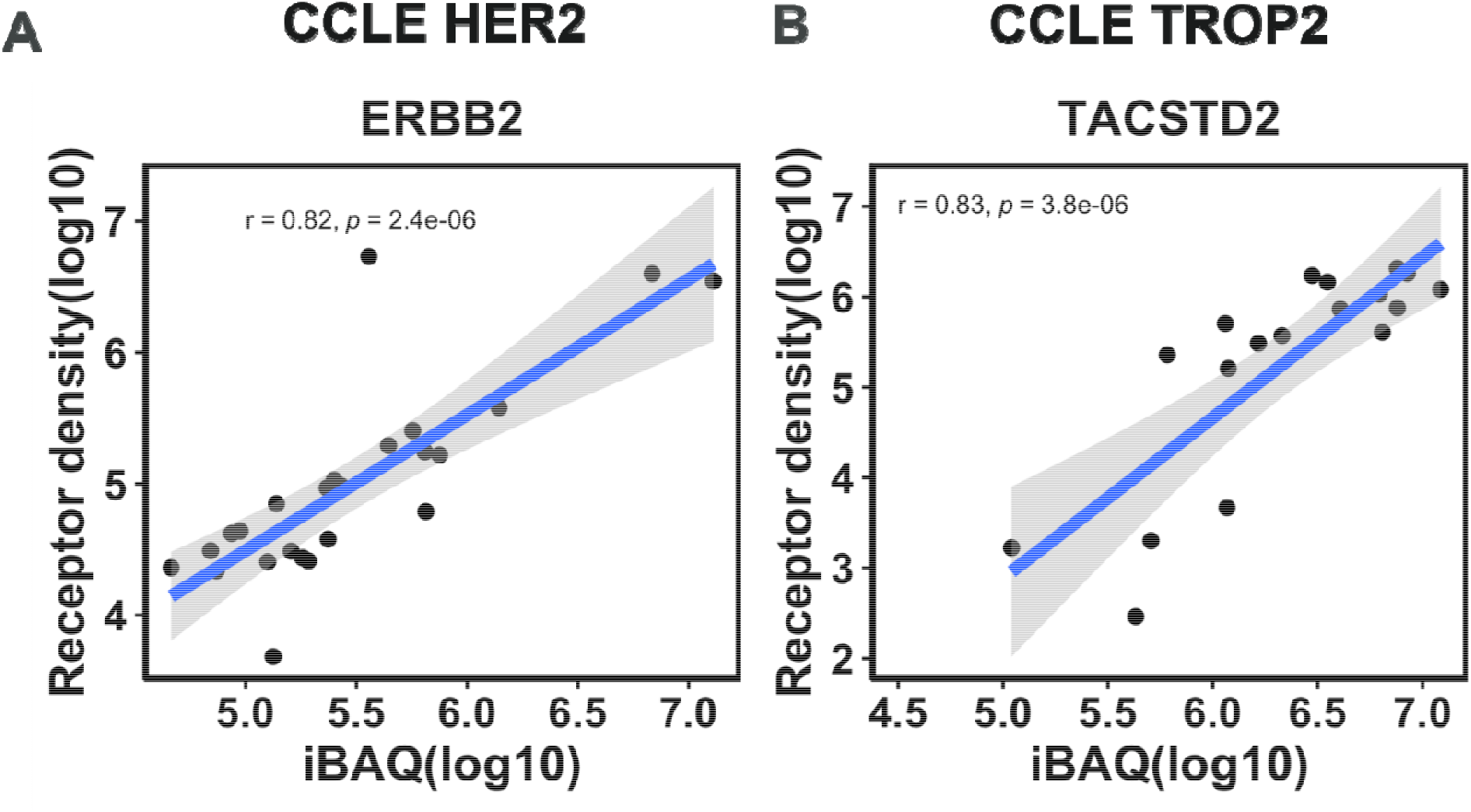
Validation of absolute protein abundance by the iBAQ method with CCLE cell line receptor density data. iBAQ quantity and receptor density for HER2 (A) and TROP2 (B) were highly correlated by Pearson correlation.

### Identifying highly expressed, tumor-enriched cell surface antigens

To identify highly expressed, tumor-enriched cell surface antigens, we ranked the upregulated and predicted cell surface proteins by their iBAQ values (Supplemental Table S2). The top five ranked antigens based on differential expression analysis and iBAQ ranking are listed in Table 1. For COAD, we identified 236 upregulated proteins at the threshold of FDR-adjusted *P* < 0.01 and fold change > 1.5. Of these 236 proteins, 46 were predicted to be expressed on the cell surface, having a SPC score of ≥1. We also evaluated well-known tumor-enriched and highly expressed cell surface antigen HER2 in BRCA. ERBB2 was highly expressed in the HER2 subtype, and iBAQ values were higher in tumor samples than in normal samples (Supplemental Figure S5), confirming that it is a highly expressed cell surface antigen in breast cancer.

## DISCUSSION

We built a comprehensive dataset containing differentially expressed tumor cell surface antigens ranked by their absolute protein expression across 10 tumor types via a newly developed computational approach (Figure 1), which represents the first surfaceome dataset containing absolute protein expression quantification derived from mass spectrometry–based proteomics data. With the addition of proteomics and phosphoproteomics characterization, CPTAC has provided the cancer community with an unprecedented view of cancer disease biology at the protein level. By combining genetic and transcriptomics data, cancer researchers have unveiled many novel mechanisms of cancer, discovered novel subtypes, and provided richer disease context for current treatments (50, 51). However, the process and parameter settings are generally not the same across tumor types in the CPTAC dataset (51) and the protein expression values are relative measurements, thus presenting challenges for studies that require cross-indication and cross-protein comparisons. Reprocessing of TMT global proteomics data with the same pipeline is necessary to avoid potential confounding factors. In addition, developing a method for estimating the absolute protein quantitation would allow cross-protein comparisons. The ability to compare and rank protein expression levels is necessary for prioritizing cancer surface antigens for novel biologics-based target identification.

To address the aforementioned challenges, we reprocessed CPTAC data sets across 10 indications with a common pipeline, FragPipe. We also incorporated missing value evaluation, implemented missing value imputation, and used a model-based method (DEP) for differential analysis, taking advantage of these recently developed proteomics data analysis methods. Through this analysis, we identified a significant number (28–417) of tumor-overexpressed proteins across 10 indications (Table 1), providing a comprehensive view of proteins that could be related to tumor pathogenesis obtained through a consistent methodology.

Although several methods have been developed and validated for absolute protein quantification from label-free proteomics data (19, 22, 52–54), there is a lack of evidence that these methods can be used for absolute quantification of TMT proteomics data. Here, we used CPTAC COAD data containing both label-free and TMT proteomics data to show that the TMT-TPA and TMT-iBAQ methods are well correlated with label-free–based LFQ-TPA and LFQ-iBAQ approaches. This finding provides a strong rationale for ranking protein expression levels using iBAQ quantification, which will ultimately aid in the identification of tumor-enriched and highly expressed cell surface antigens. To further evaluate our approach of using TMT-iBAQ for estimating absolute protein quantification, we reprocessed CCLE TMT proteomics data using the same pipeline and calculated iBAQ values for all identified proteins. Our analysis revealed a high correlation between iBAQ values and receptor density of HER2 and TROP2 in about 30 selected breast and lung cancer cell lines (26, 55), providing additional confirmation for using the iBAQ absolute quantitation method for TMT proteomics data.

A limitation of our study is that we tested our approach using only HER2 and TROP2 receptor density data in a limited number of cell lines. Additional orthogonal measurements of protein absolute quantification for a larger number of proteins will help to further validate the iBAQ methodology for the estimation of absolute protein abundance using TMT proteomics data.

In summary, we have established a novel computational approach for identifying tumor-selective and highly expressed surface antigens from proteomics data, thereby providing a unique data resource for integrating potential cancer disease vulnerabilities employing biologics-based cancer therapeutics. Our results demonstrate that differential protein expression analysis combined with iBAQ ranking can aid in identifying tumor-enriched and highly expressed cell surface antigens. We evaluated the TMT-iBAQ and TMT-TPA approaches with label-free data from the CPTAC COAD data set and found that TPA and iBAQ could be applied to TMT proteomics data to derive estimated absolute protein abundance. We further validated the iBAQ protein abundance approach with receptor density data from the CCLE cell line. Collectively, the data shared in this report provide a method for the identification of tumor-enriched and highly expressed cell surface antigens with iBAQ and DEP analysis. FragPipe-derived DEP, TMT-TPA, iBAQ, and iBAQ-derived copy numbers are useful data sources for cancer biology research and the identification of potential targets. The methodology and the data sources in this study can serve as valuable resources for cancer research.

## Data Availability

The FragPipe output of protein abundance estimation, derived TMT-TPA, iBAQ, iBAQ-derived copy number and differential protein expression data for CPTAC ten indications can be downloaded from https://doi.org/10.5281/zenodo.7991979.

## Supplemental data

This article contains supplemental data.

## Author Contributions

J. W. and W. Z. conception and design; A. P., M. W., and M. S. flow experiment; J. W., Y. W., R. D., J. B., E. H., G. D., B. S., Z. L., and W. Z. analysis and interpretation of data. All authors contributed to the writing, review, and/or revision of the manuscript, have approved the final version of the manuscript, and agree to be accountable for all aspects of the work.

## Acknowledgments

We gratefully acknowledge the CPTAC for providing open-source proteomics data and Deborah Shuman of AstraZeneca for editing the manuscript and formatting the figures.

## Abbreviations

The abbreviations used are:

APEX: absolute protein expression
BRCA: breast cancer
CCLE: Cancer Cell Line Encyclopedia
ccRCC: clear-cell renal cell carcinoma
COAD: colon adenocarcinoma
CPTAC: Clinical Proteomic Tumor Analysis Consortium
DEP: differential protein analysis
DIA: data-independent acquisition
FDR: false discovery rate
GBM: glioblastoma multiforme
HNSCC: head and neck squamous-cell carcinoma
iBAQ: intensity-based absolute quantification
IgG: immunoglobulin G
KNN: k–nearest neighbor
LFQ: label-free quantification
LSCC: lung squamous-cell carcinoma
LUAD: lung adenocarcinoma
NAT: normal adjacent tissue
OV: ovarian cancer
PBS: phosphate-buffered saline
PDA: pancreatic ductal adenocarcinoma
SPC: surface prediction consensus
T-DXd: trastuzumab deruxtecan
TMT: tandem mass tag
TPA: total protein approach
UCEC: uterine corpus endometrial carcinoma.

