## Supplemental data for "Pan-cancer Proteomics Analysis to Identify Tumor-Enriched and Highly Expressed Cell Surface Antigens as Potential Targets for Cancer Therapeutics"

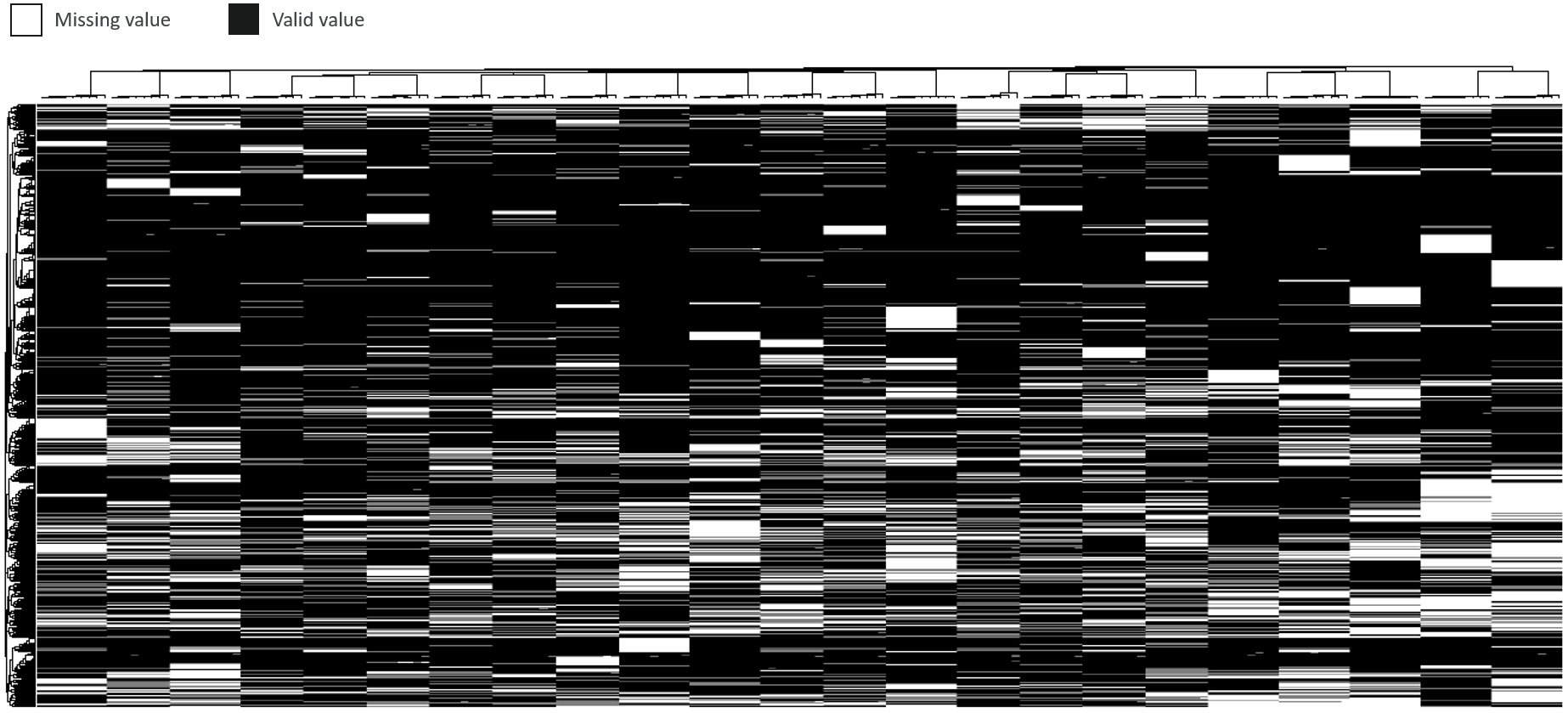

**Figure S1.** Missingness pattern in data from the Clinical Proteomic Tumor Analysis Consortium (CPTAC) for the clear-cell renal carcinoma indication. Missing data showed missing-at-random pattern across tumor and normal conditions. Multiplex-level missing pattern is missing not at random.

CPTAC COAD

LFQ

TMT 10-plex

MaxQuant

FragPipe

DDA-TPA, iBAQ

100 tumor samples

2597 proteins

97 tumor samples

9709 proteins

Abundance ratio

93 matched tumor samples

TMT-TPA, iBAQ

**Figure S2.** CPTAC colon adenocarcinoma (COAD) data set and workflow. Label-free proteomics raw intensity data of colon cancer from the CPTAC were processed with MaxQuant. Tandem mass tag (TMT) 10-plex mzML files were processed with FragPipe. Label-free raw intensity and TMT abundance data were analyzed by the total protein approach (TPA), and intensity-based absolute quantification (iBAQ) was used to derive estimated absolute protein abundance. DDA, data-dependent acquisition; LFAQ, label-free absolute protein quantification.

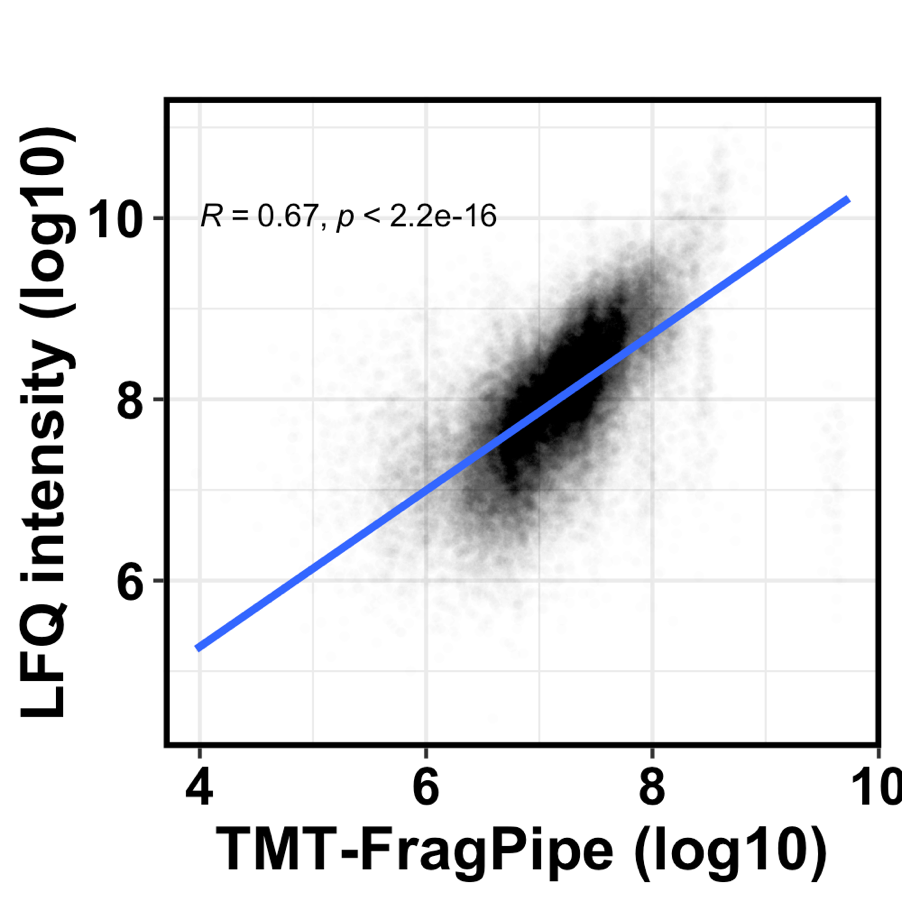

**Figure S3.** Correlation between FragPipe TMT-based protein abundance and label-free quantification (LFQ) intensity.

**A**

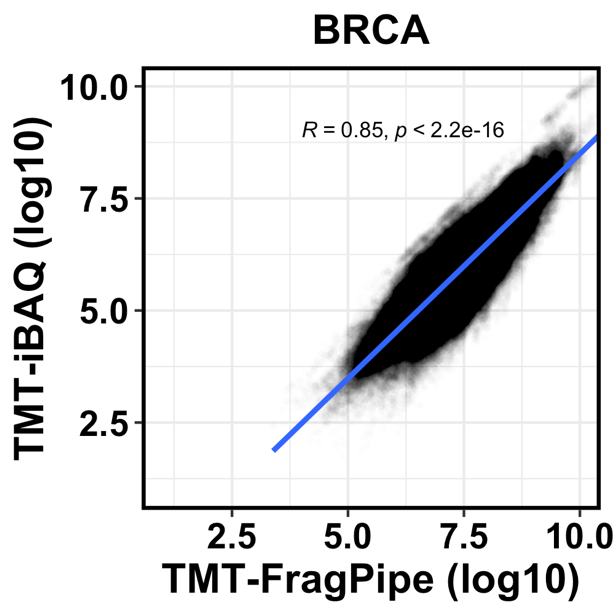

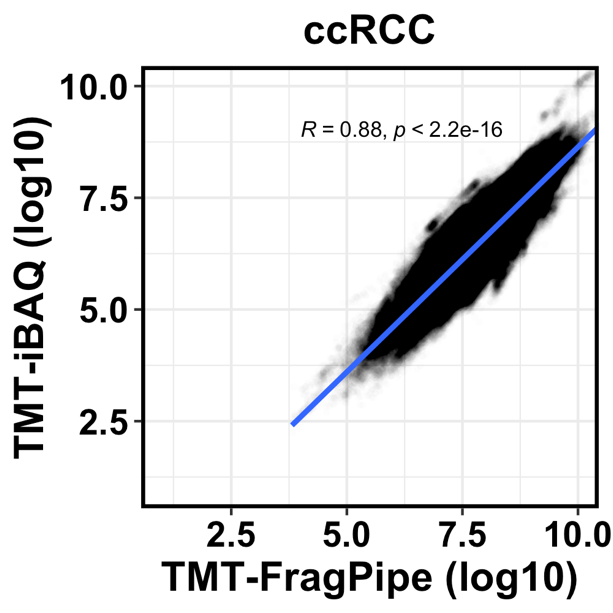

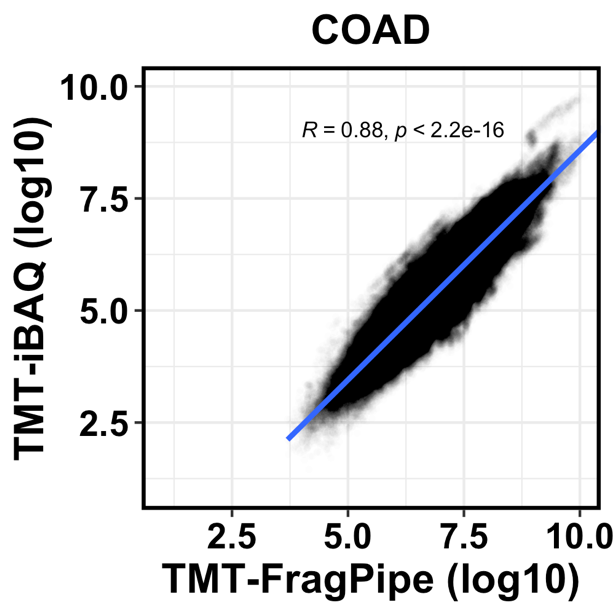

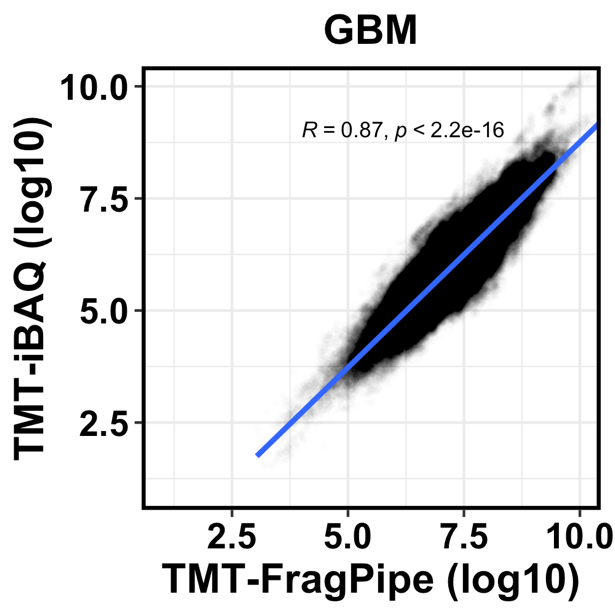

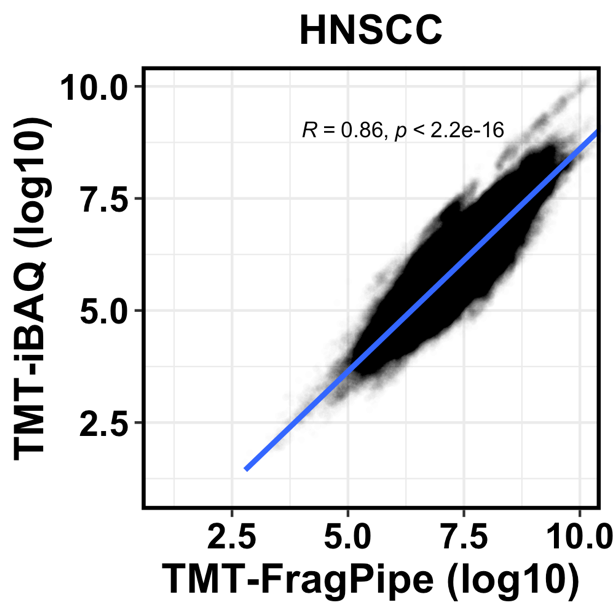

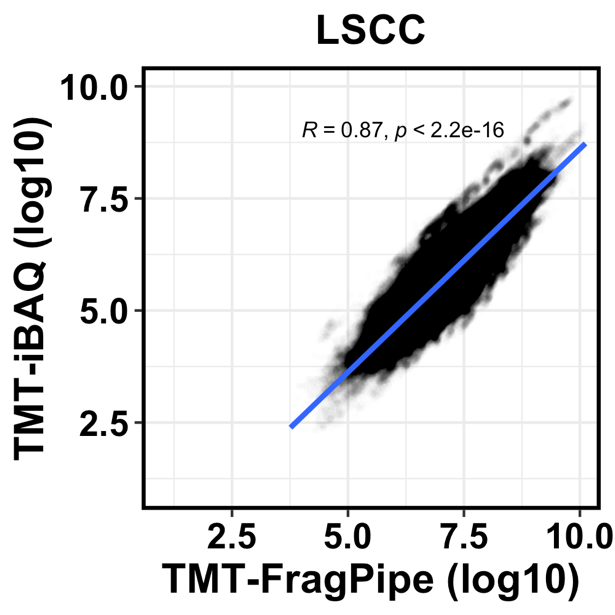

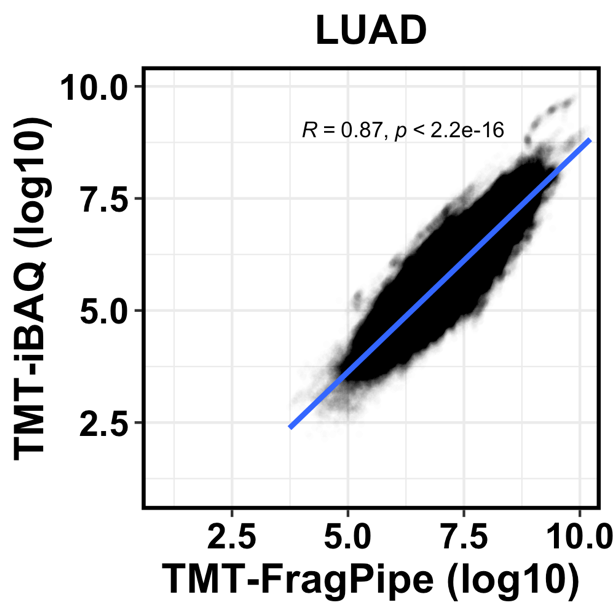

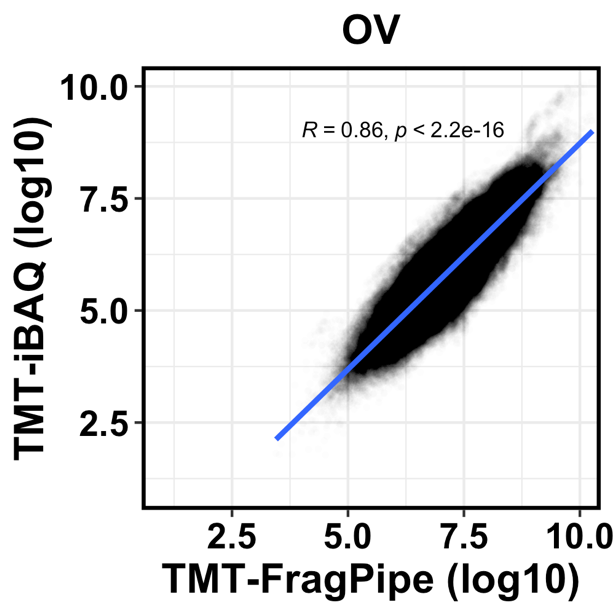

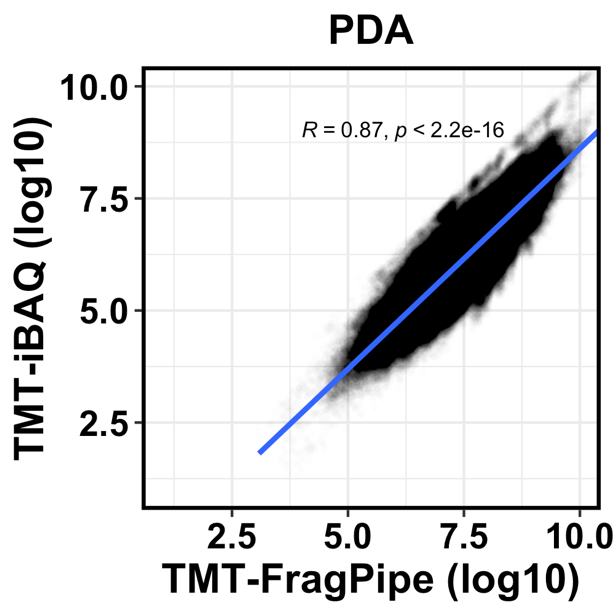

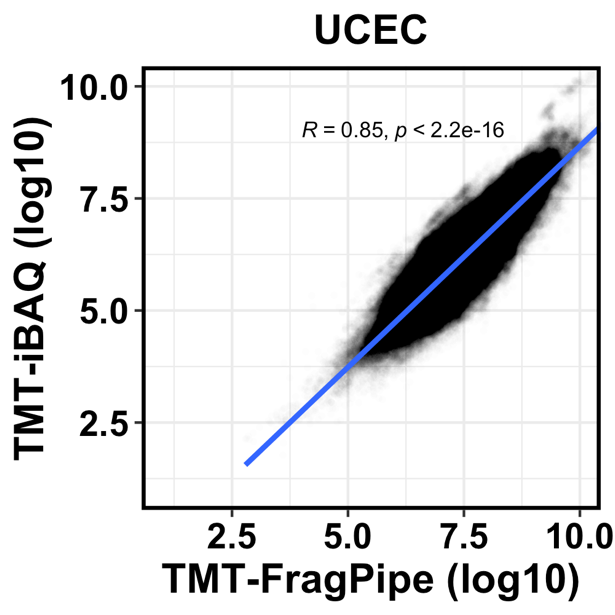

**Figure S4.** The correlation between protein abundance reported by FragPipe and derived IBAQ for CPTAC.

**A**

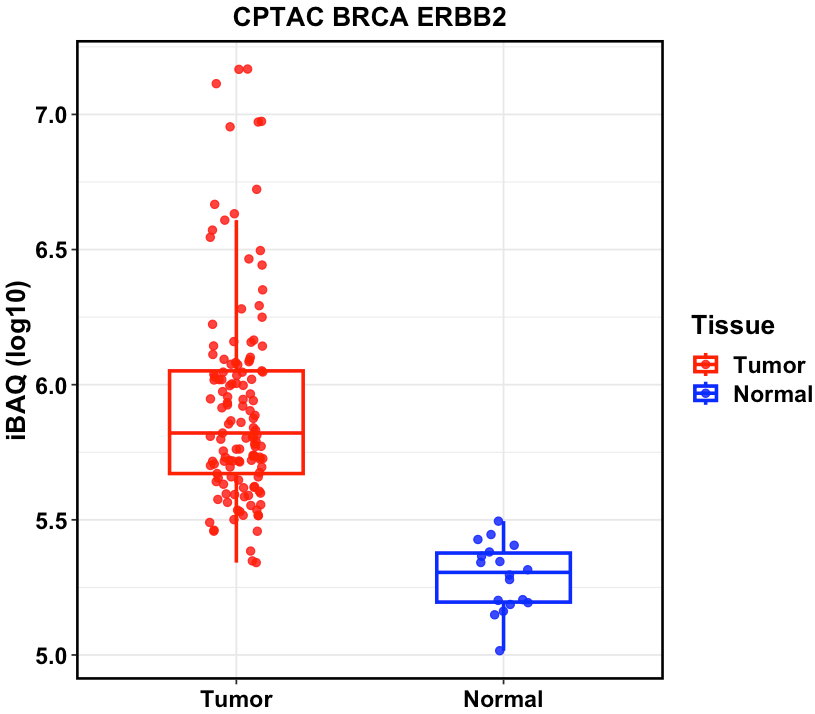

**B**

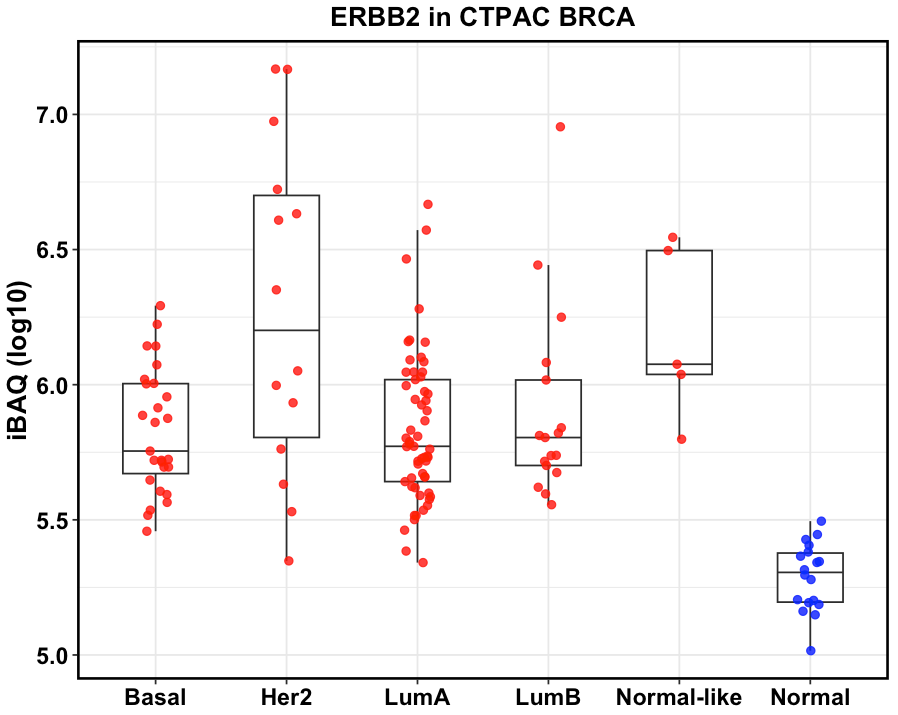

**Figure S5.** Case study in CPTAC breast cancer (BRCA) indication. (A). iBAQ quantity could identify ERBB2 as highly expressed cell surface antigen as high iBAQ in tumor compared with normal samples. (B). Her2 expression is higher in HER2 subtype in CPTAC BRCA.
